# Low cost conservation: Fishing gear threats to marine species

**DOI:** 10.1101/2020.08.30.273532

**Authors:** Tim Cashion, Travis C. Tai, Vicky W.Y. Lam, Daniel Pauly, U. Rashid Sumaila

**Affiliations:** Fisheries Economics Research Unit, Institute for the Oceans and Fisheries, University of British Columbia; Changing Oceans Research Unit, Institute for the Oceans and Fisheries, University of British Columbia; *Sea Around Us*, Institute for the Oceans and Fisheries, University of British Columbia; School of Public Policy and Global Affairs, University of British Columbia

## Abstract

Understanding conflicts between objectives of fisheries and conservation is the key to finding win-win situations for marine biodiversity and fishers. Many marine species are threatened by harmful interactions with fisheries, but the threats they face are associated with the fishing gear used. Here, we undertake a novel analysis of marine species and their gear-specific threats to evaluate conservation-fisheries trade-offs to identify areas with high competing goals. Our analysis suggests that gillnet and longline fisheries pose the greatest risk to marine species yet deliver relatively low profits, emphasizing the inefficiencies of these gears. We find that the majority of the high seas has low economic fisheries benefits with over 25% of the high seas categorized as areas of ‘conservation prioritisation’ over fisheries.

## Introduction

Fishing is a major threat to many marine species globally (*1*). However, different fishing gear types are heterogeneous in their spatial extent and their impacts on different species. Therefore, treating different fisheries homogenously with regard to their spatial management likely causes unnecessary conflict between fisheries and conservation priorities, and may impose too high costs without necessarily achieving their intended conservation goals. Here, we aim to understand the threats due to specific fishing gear types and their spatial overlap with species of conservation concern. Recent studies have highlighted the spatial extent of fisheries (*2*, *3*), as well as the fisheries risk to species of conservation concern (*4*). While fisheries pose obvious risks to both targeted and bycatch species (*5*), they also provide livelihoods and food security to millions and billions, respectively (*6*). Thus, the goals of fisheries and conservation ought to be balanced to minimize the costs and maximize the benefits where possible.

Trade-offs, such as those between fisheries and conservation, are increasingly important to consider and analyze for marine spatial planning as multiple sectors need to be taken into account. For example, White et al. (*7*) evaluated the trade-offs necessary when implementing ocean-based wind turbines that limit access to others sectors such as tourism (e.g., whale watching) and fisheries. However, when considered as a whole, the placement of these turbines can be done in a way that increases the value of the area as a whole while minimizing losses for certain sectors (*7*).

Previous analyses have demonstrated these trade-offs between fisheries and ecosystem health (*8*). However, win-win situations are often shown to occur in overly simplified models that do not account for all variables such as employment for fishers (*8*) or spillover impacts to other areas of importance (*9*). It is therefore important to consider the different scales at which these conservation plans act and the implicit trade-offs between social and ecological outcomes in many fisheries management plans (*9*). In addition, taking the heterogeneity of fisheries into account can lead to more positive fishery outcomes without compromising conservation goals (*10*).

Trade-off analysis is especially important in areas where the units are not directly comparable. Protecting species of conservation concern from their fisheries threats is one such case where fisheries are valuable for their contribution to livelihoods and food security (*11*), while protecting marine biodiversity is important for its contribution to various ecosystem services including fisheries production, tourism, and other regulating services (*12*, *13*). These trade-offs can often be managed through marine spatial planning or other forms of spatial management. Ideally, this can allow different sectors to thrive in their optimal areas while restricting them from operating in the optimal areas for other sectors, which is necessary for managing conflicts between different stakeholders and resource users.

Here, we establish the first estimate of large-scale conservation trade-offs when protecting species from their major fishing gear threats. This analysis is based on 4,579 marine species included in the International Union for the Conservation of Nature (IUCN) Red List with *specific* fishing gear listed as threats. We use species’ distribution range maps with their threat status (*14*) (for the 2,226 out of 4,579 with distribution maps and gear threats) and combine them with a spatialized fisheries catch by gear database (*15*). We use gear threats by species described by the IUCN, and weight IUCN conservation status on a linear scale to adapt the biodiversity risk score (*16*) as an ‘average’ threat status of marine species within an area *by fishing gear*. We also use a weighted threat score based on these same criteria but not scaled by the number of species for all marine areas of the world (Table S1). These two metrics evaluate the average threat status of marine species (biodiversity risk score) and the total threat to marine species (weighted threat score) in a given area. We then combine this with fisheries catch and profit data by gear type to highlight areas with low-cost trade-offs between fisheries and conservation, and areas of high conservation concern that are also highly important to fisheries where there is likely to be competition between fisheries and conservation objectives.

## Materials and Methods

### IUCN data

The International Union for Conservation of Nature (IUCN) maintains the IUCN Red List of Threatened Species (hereafter, ‘Red List’) that documents the population status and threats of species globally. Species (and sub-populations of species) that are assessed by the IUCN are categorised into one of the following in order of increasing conservation threat: Least Concern, Near Threatened, Vulnerable, Endangered, Critically Endangered, and Extinct. A final category exists of ‘Data Deficient’ that indicates there is not enough information to properly assess the population status of a species. Together, Vulnerable, Endangered, and Critically Endangered are often grouped together as ‘Threatened’. In addition to the categorisation of the threat status, the IUCN provides species range maps, detailed description of threats, and other important information on the species included in the Red List.

Red List Categories, spatial habitat maps, identified threats by species, and a description of threats were gathered from the IUCN API version 2019-2 (*14*). The known threats to each marine species were extracted from the IUCN API focusing on threats identified in Category 5: Biological Resource Use. The Red List identifies four different types of fisheries impacts from either intentional or unintentional capture/harvest from the large- or small-scale fisheries. These threats are categorised by the IUCN as unknown, past, no, low, medium, or high impact. In addition, the text from the ‘Detailed Threats’ and ‘Use and Trade’ sections for each marine species was extracted from the IUCN API. After extracting the description of the species threats and use and trade information, we tokenized (i.e., separated the text into one and two word strings) the text extracting standard stop words and searched for single words and bigrams of fishing gear types (e.g., ‘bottom trawl’ or ‘longline’, see Table S2). Bigrams were used as they can include more specific gear types than single words alone (e.g., ‘bottom trawl’ versus ‘trawl’ and ‘purse seine’ versus ‘seine’). We use the presence of these words within these narrative sections as a proxy for a particular gear being a threat to these species. The labels for different fishing gear types analysed are included in Table S2. These were devised to match the IUCN narrative text most closely to relevant *Sea Around Us* gears (Table S4).

We accessed spatial species distribution shape files from the Red List for comprehensively assessed groups that include marine organisms (*14*). We supplemented this with Bird Life International species distribution files for birds that occur in marine areas (*37*). Marine molluscs have not been comprehensively assessed (i.e., less than 80% of the species in this group have not been assessed) by the IUCN except for Cone Snails (*Conus* spp.), and this is a gap in our data coverage. We converted the species distribution maps to raster format at various spatial scales to correspond to existing datasets in fisheries research, but used a 0.5° latitude by 0.5° longitude for our analysis as it matches the *Sea Around Us* fisheries catch database. We rasterised the species distribution files following the same process of O’Hara (*16*).

We used the same prioritisation of species IUCN Categories based on regional assessments where possible as in O’Hara (*16*). Here, regional assessments were matched to associated (i.e., overlapping) marine ecoregions (*38*) that were then associated with their corresponding cells of the spatial grid used. Regional assessments were used in preference of global assessments where possible.

The Red List contains detailed information on the species system and habitat types. The system assigns species to terrestrial, freshwater, and marine ecosystems (with species able to occur in multiple ecosystems) and this was used to restrict our analysis to marine species. In addition, we restricted those within the marine ecosystem to specific habitats to eliminate species that are not dependent on the marine environment nor likely to be affected by fisheries, following O’Hara et al. (*16*). After restricting our analysis to marine dependent species that are threatened by fishing gear and have species range maps available from the IUCN or BirdLife International, our analysis focused on 2,226 number of species (Fig S2).

### Fisheries data

The *Sea Around Us* has reconstructed marine fisheries catches for all fishing countries and territories from 1950 to present. The process of reconstruction supplements and corrects reported fisheries catches with estimates of known to be overwhelmingly excluded fishing sectors (e.g., artisanal and recreational fishing) and practices (e.g., discarding) (*39*). These catches are spatially allocated according to a rule-based assignment and based on known information on where the fishery was operating (within domestic EEZs, foreign EEZs, the high seas, etc.). The spatial scale used is 0.5° latitude by 0.5° longitude.

The reconstructed catches were then assigned to their respective fishing gear types for industrial and small-scale sectors. This process relied on similar methods as the catch reconstructions relying on official catch statistics by gear type, as well as fisheries reports, catch surveys, newspaper articles, and other grey literature. The process by which each countries fishing gear was assigned is documented in Cashion (*18*). The result is a catch database with gear information included for all catches.

We used the *Sea Around Us* database (v.47) for catches by gear type (*15*, *18*) and were accessed from the *Sea Around Us* database by the first author. This database includes spatially allocated reconstructed fisheries catches by gear type and taxon for all fishing countries and territories of the world. The spatial scale of this dataset is 0.5° latitude by 0.5° longitude and has been used in many global fisheries studies (*22*, *40*).

In addition, we use the corollary Fisheries Economics Research Unit (FERU) ex-vessel price database that includes reported and estimated first-sale prices for all taxa for each fishing country by year (*41*) to incorporate the potential lost revenues for fisheries closures in areas of high biological importance. Revenue is calculated as the ex-vessel price (real 2010 USD per metric tonne) of a specific taxon caught by a country in a given year multiplied by its landings amount in metric tonnes.

Due to the importance of gear to our analysis, and its importance as a determinant on the ex-vessel price of fish (*42*–*44*), we modify the ex-vessel prices by a gear multiplier. We determine this gear multiplier through a hedonic pricing model where gear type is an explanatory variable of the ex-vessel fish price.

We used the U.S. National Marine Fisheries Service annual commercial landings by gear type (*45*). We harmonized the gear types listed to match our existing dataset gear types. While this dataset is not representative of global fisheries nor ex-vessel prices for all species, it gives adequate coverage to derive the effect of different gear types used for catching different species and how it modifies ex-vessel prices.

We then used a fixed-effects model with linear regression to derive the effect of changing gear type on the ex-vessel price *Price_xyzt_* for species *x*, gear *y*, in year *t*. We used the natural logarithm of landings and prices as these variables are closer to a normal distribution when log-transformed, and thus reduces potential heteroscedasticity in our residuals. We also use the country, species, and year as explanatory variables to account for other changes in the price both over time and between these different markets.

Our regression equation is:

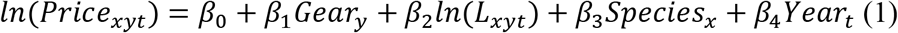

We used the estimated gear-type coefficients as gear specific multipliers *β*_1_*y*__ that will modify the ex-vessel price of fish caught (Table S5). The multiplier values for each gear type range from 0.54 to 1.34.

Finally, we used an updated version of FERU’s cost of fishing database (*46*) to account for the cost of fishing that varies by country and gear type. Cost of fishing is broken down into its component parts in the database (e.g., fuel, labour, capital, maintenance, etc.), and here we use the total cost by gear type and country per tonne of fish landed (*C_yz_*). Where the cost of fishing was not available for a particular gear and country, we used the regional average for that gear type, and where this was not possible we used the average cost across gear types for the region. We used the FAO socioeconomic regions for this stage of the analysis (*47*). We then derive the profit by gear type and country for each cell *i*, based on the catch by gear type multiplied by the ex-vessel price multiplied by the gear type multiplier (*M_y_*) minus the cost of fishing for that amount of landings. Therefore, profit (*π*) in cell *i* is equal to:

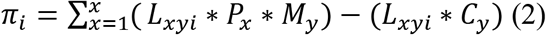

Both catch and profits are expressed throughout in tonnes/km^2^ and $/km^2^ calculated based on the water area in each 0.5° by 0.5° cell. We would expect profit to be relatively equal across gear types, as fishers would switch production systems if one was seen as more profitable. However, profit is expected to vary by gear type in practice in regulated fisheries due to restrictions on switching gear types or target species in regulated limited-entry fisheries (i.e., limited license or quota availability). Further, it is likely that it varies in fisheries not regulated by limited entry through high transition costs of switching gear types, which can be further restricted by credit constraints, and distortions from subsidies which are not equal among fleets (*48*).

Our analysis focuses on the trade-offs among different fisheries with specific gear types. As such, we did not use the Global Fishing Watch data where gear types are often aggregated (bottom and pelagic trawlers grouped together as trawlers), or only represent a narrow subset of the fleet (e.g., drifting longlines instead of all longlines) (*2*). While the Global Fishing Watch data covers a large part of the global fishing effort in non-coastal areas (between 50% and 70% in areas greater than 100 nautical miles from land (*2*)), it does not have the same coverage of vessels. The dataset is biased against small-scale vessels which are not harm-free.

### Analysis

First, we analysed the major fishing gear threats based on their appearance in the narrative text of the Red List species profiles. We associated these descriptors to their gear types and examined the number of species by Red List Category and weighted threat status by gear type. We used a linearised weighted scoring method to weight the presence of a species by its Red List Category (e.g., ‘Least Concern’ = 0, ‘Endangered’=0.6, etc., Table S1) (*16*). Older and outdated categories were updated to their current descriptors such as ‘Lower Risk/near threatened’ to ‘Near Threatened’.

We mapped areas of high fisheries interest and high conservation concern (Figure S1). The areas with high overlap between conservation concern but low fisheries catches and effort are areas that could be fully protected from these threats for the long term (termed here, ‘conservation prioritisation’). However, the areas with high overlap of fisheries interest and high conservation concern are where the impacts on species may be greatest both in terms of fisheries threats and the potential benefits for conservation from fisheries closures. We used this categorisation into four quadrants to simply but effectively delineate areas between their contribution to fisheries and their risk to species of conservation concern from fisheries.

We use three main metrics frequently in our analysis. First, we adapted the biodiversity risk score developed by O’Hara et al. (*16*). This indicator is a measure of the average conservation threat status of an area of the marine environment (Equation 3). It is expressed as:

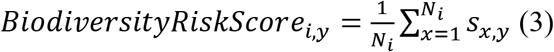

Where *N* is number of assessed species in the cell *i*, *s_i_* is the linearized Red List Category numeric score (*s*) of species (*x*) threatened by fishing gear (*y*). All biodiversity risk scores are thus values of between 0 and 1 that roughly correspond to the linear Red List Category scale described above (Table S1). Our adaption of this metric limits it to species threatened by a specific fishing gear (*y*).

Second, we adapt this metric so that it is not normalised to the mean threat in the cell but representative of scale (number of species) and severity (Red List Category) of threat. This metric is named the ‘weighted threat score’ defined as the sum of species present multiplied by their linearized Red List Category numeric score. We do not normalise (divide by the number of species in a given cell) this to highlight areas of conservation concern based on the number of species in that cell in addition to their threat status.

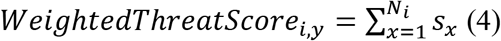

This analysis was conducted in R (*49*), and used the tidyverse (*50*), rMarkdown (*51*), tidytext (*52*); gridExtra (*53*); sf (*54*), rgdal (*55*), cowplot (*56*), and wesanderson (*57*) packages. All code and outputs are available at www.github.com/timcashion/iucnfishingthreats.

## Results

### Fishing threats to IUCN species

For the 14,126 marine species included on the IUCN Red List, fishing is identified by the IUCN as a threat to 4,455 of them (31.5%) (Threat category 5.4: fishing and harvesting aquatic resources). This threat is from both large- and small-scale sectors, and from intentional and unintentional capture (i.e., by-catch, Figure 1A). According to the identified threats, the small-scale sector is a threat to a greater number of species than intentional or unintentional capture by the large-scale sector. Interestingly, the impacts of fishing are identified to be low or unknown on most of the species, whereas medium or high fishing impacts are identified for only very few species.

**Fig. 1.**
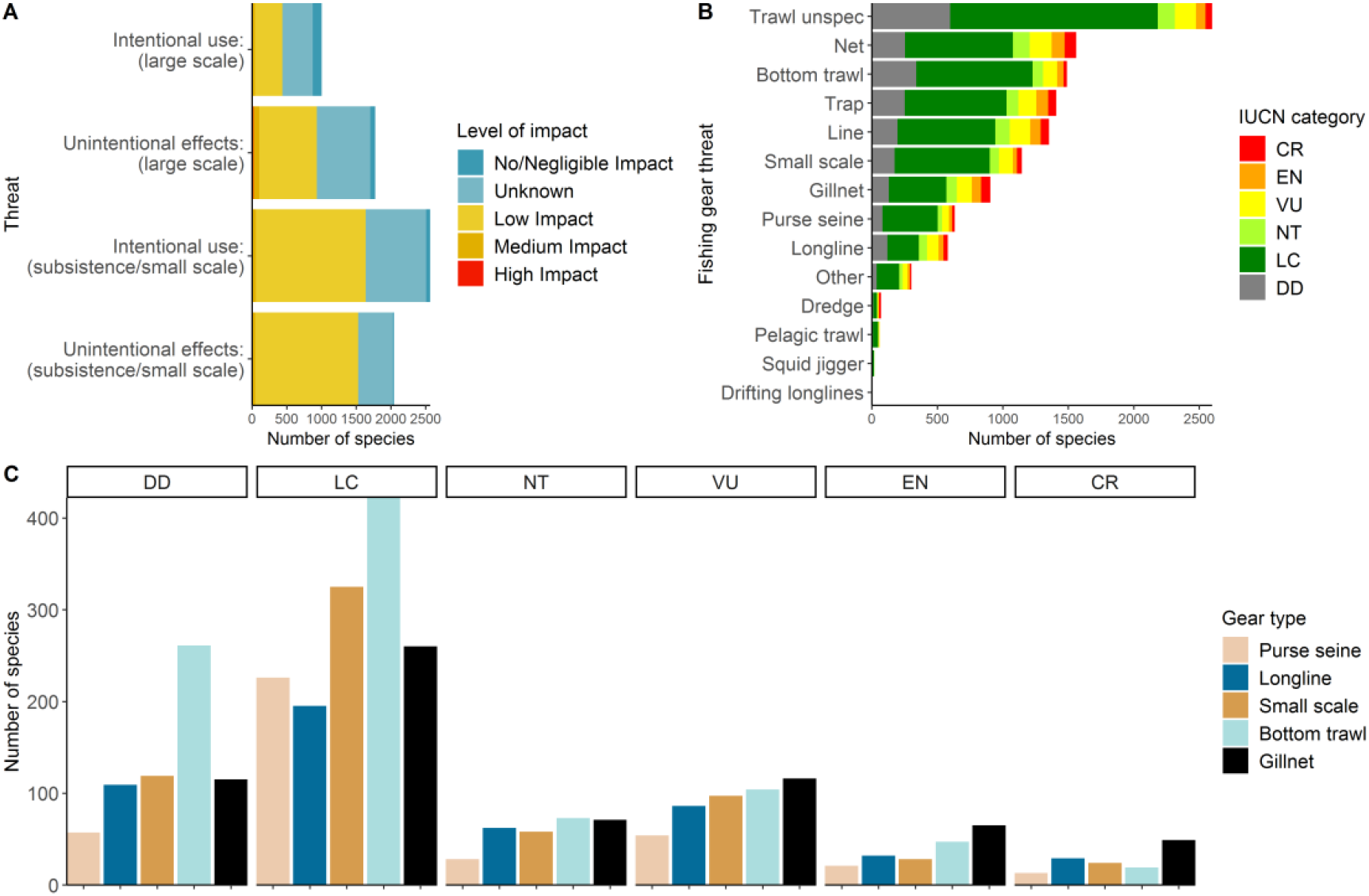
IUCN fishing gear threats: species categorized by fishing gear threat (A); fishing gear threats by gear types listed (B); and identified gear types by threat status (C). Note: DD: Data Deficient; LC: Least Concern; NT: Near Threatened; VU: Vulnerable; EN: Endangered; CR: Critically Endangered.

Within the Red List, trawls are identified as a gear threat for more than 2,500 marine species (Figure 1B). While many of the species caught by this gear type have an elevated risk of extinction (Near Threatened and higher), most are either Least Concern or Data Deficient within these gear types. The ‘gear types’ that appear in most species threat description are generally more vague gear terms (e.g., ‘trawls’ and ‘nets’) and could be attributable to several types of fishing gear (e.g., trawl nets, gillnets, seine nets all fall under ‘nets’). The more specific gear categories identified are used in the remaining parts of the analysis (Table S2).

From the gear types we have previously identified (Figure 1C) we see again that bottom trawl fisheries are associated with the largest number of Red List species. However, we see these species are mainly in the Data Deficient, Least Concern, and Near Threatened categories while gillnet gears have the most number of species in the Threatened categories (Vulnerable, Endangered, and Critically Endangered).

### Profits of fishing fleets

Our estimates of profits by fishing fleet vary between fleets as well as geographically (Supplemental figures). Overall, most fleets profits have a roughly normal distribution with a mean near 0 and values extending into large negative values as well as large positive values (Figure S5). While the result of some areas having negative profits may seem counterintuitive, this is to be expected given heterogeneity of fisheries, especially spatially, in addition to our measure of profits not including subsidies. This finding is also supported by economic theory where rents of open-access fisheries are expected to be 0 (*17*). Our profit results are driven by the total revenues from each gear type in each 0.5° by 0.5° cell minus the average cost of operating these gears by each country. In this way, these estimates give a valid approximation of the spatial value of fisheries benefits.

### Low-cost solutions to gear threats and conservation

Globally, annual fisheries profits by bottom trawl fisheries are estimated to be at an average of $57 real USD per km^2^ of area fished (Table 1). However, an average of 11 species are targeted by bottom trawls in each of the half degree by half degree grids cells with this gear, with an average biodiversity risk score of Near Threatened (0.19). Taken together, the generally high value of their catches make them perform relatively well when considering their average biodiversity risk score and their profits together. In contrast, longline fisheries have low profits per area occupied ($11) as they fish over a large spatial area. While they operate in areas with a similar biodiversity risk score as bottom trawl fisheries (0.19), they do so with much lower returns meaning they produce less fisheries profits for their relative conservations risk (weighted threat score) than bottom trawl fisheries. It is important to note that these profits are taken based on the estimated revenues and costs of these fishing vessels, without taking into account subsidies. Therefore, the private profits for these fisheries are higher than shown here.

**Table 1.** Mean values of measures of fisheries conservation concern and profit by gear type with standard deviations in brackets. Mean values are average of all 0.5° by 0.5° cells where that gear is present.

| Gear type | Profit (\$/km<sup>2</sup>) | Number of species | Weighted Threat Score | Biodiversity Risk Score |
| --- | --- | --- | --- | --- |
| Bottom trawl | 57.38<br>(613.94) | 10.9 (14.5) | 1.98 (2.98) | 0.19 (0.12) |
| Gillnet | 11.87<br>(134.83) | 35.4 (18) | 5.95 (3.67) | 0.16 (0.04) |
| Longline | 10.65<br>(96.44) | 25.5 (12.7) | 5.01 (2.73) | 0.19 (0.04) |
| Pelagic trawl | -125.79<br>(493.41) | 1.8 (1.9) | 0.2 (0.44) | 0.06 (0.11) |
| Purse seine | 9.19<br>(226.52) | 22 (13.7) | 3.9 (2.61) | 0.16 (0.06) |
| Small scale | 548.91<br>(3409.66) | 23 (28.3) | 2.81 (3.28) | 0.13 (0.06) |

We divided the global ocean into half degree by half degree grid cells. If we consider each cell in the grid as independently managed, we can examine the trade-offs in each cell based on its fisheries profits and conservation value (Figure 2). Although bottom trawl fisheries operate in many spatial cells of conservation concern (high ‘weighted threat score’ values), they also generate substantial profits from these fisheries. Gillnets, alternatively, have a large number of cells that are below the median value for fisheries profits and have high weighted threat scores. Pelagic trawls have low weighted threat scores overall and thus the reduction in their use may not lead to large conservation gains. However, pelagic trawls are shown to be non-profitable ($-126) and it means this gear type is inefficient. Hence, persistent use of this gear type may not be economically beneficial to human well-being.

**Fig. 2.**
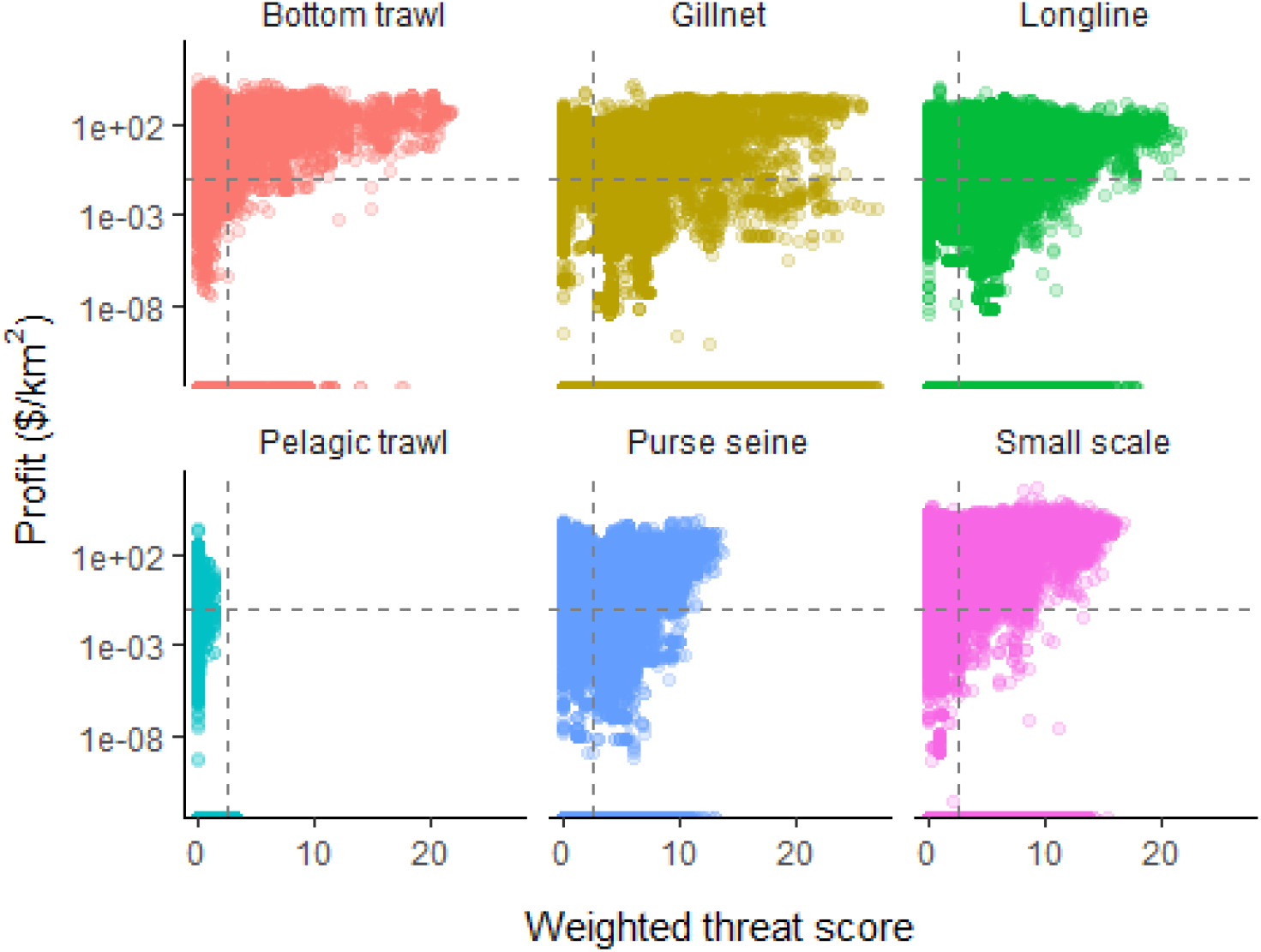
Fisheries profits (log scaled) and weighted score for major gear types. Each point represents a 0.5° by 0.5° cell. Grey dashed lines indicate the median values (excluding values of 0) across gear types for fisheries profits per km^2^ and weighted threat score.

These results by gear type also show spatial variability. Longline gears are used for a wide diversity of species and their spatial profitability varies over the globe (Figure 3A). In addition, their weighted threat score shows hotspots in the Mediterranean and the waters surrounding Indonesia with links between continents from trans-oceanic species such as the oceanic whitetip shark (*Carcharhinus longimanus*) (Figure 3B). The categorization shows large areas of the ocean where fisheries and conservation are both important thus placing them in the ‘competition’ quadrant but many EEZs are categorized as ‘conservation prioritization’ and with the polar areas generally being ‘areas of low concern’ for this gear type (Figure 3C). These results provide a broad view of the trade-offs of protecting species of conservation concern from their gear-specific fisheries threats and the monetary benefits of these fisheries. Broad overviews of conservation-fisheries tradeoffs can guide marine spatial planning efforts, especially in areas with overlapping fishing (and non-fishing) activities. Competition areas and areas closer to the median of the tradeoff analysis (Figure 2), will require greater effort and compromise by stakeholders to balance trade-offs. Similar figures for other gear types are available in the supplemental materials (Figure S6-S9).

**Fig. 3.**
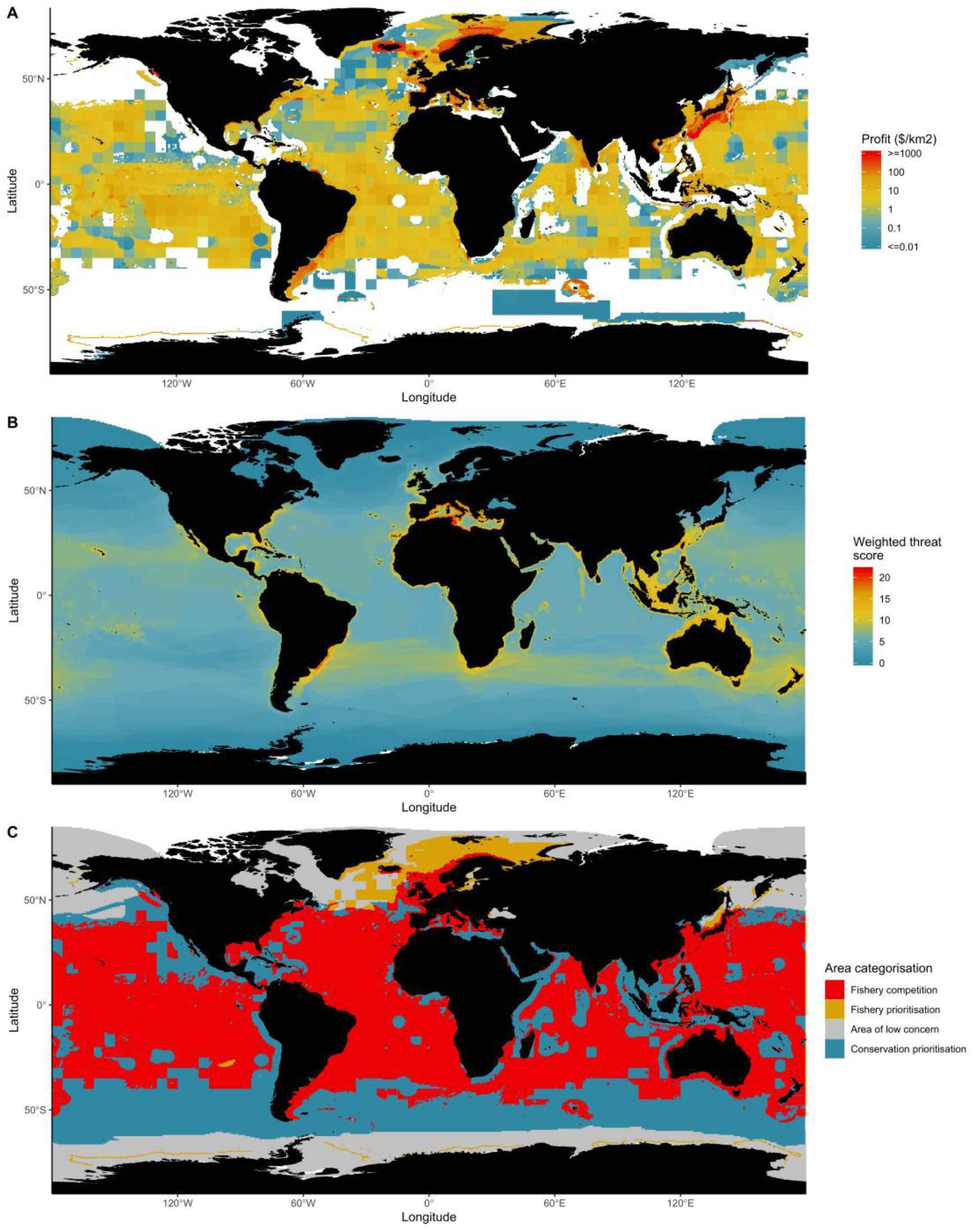
Longline: Distribution of profits (A), weighted threat score (B), and categorisation (C).

The six major gear types included here (Table 1) together account for 92% of global fisheries catches (landings and discards) (*18*). Bottom trawls, gillnets, and longlines are used in large extent (i.e., have many more cells) where the weighted threat score is higher than the median, especially in contrast to pelagic trawls, purse seines and small-scale gears. Among these higher impact gears, the fisheries profit gained in each grid cell varies substantially. Interestingly, gillnet fisheries are operating in many areas of high conservation concern with fisheries profits below the median value (across gear types). This demonstrates that the social costs of this gear are higher than previously thought (*19*), and the social benefit may not be net positive given the relatively low profits achieved (mainly below the median). In contrast, bottom trawl fisheries while overlapping on the weighted threat score dimension with gillnet fisheries, achieve higher profits thus representing a conflict between conservation goals and fisheries goals. Small-scale gears and purse seine have mixed results with weighted threat scores not nearly as high as gillnet or bottom trawls, and a mix of high and low profit areas. Pelagic trawls are often used solely for relatively low-value species (from krill to Alaska pollock), but have very low weighted threat scores throughout their range of fisheries profits.

### Protecting the high seas

Areas beyond national jurisdiction (i.e., the high seas) have recently received increased attention for their protection for biodiversity and fishery gains (*20*, *21*), while leading to little losses in terms of food security (*22*, *23*). According to our framework, the high seas have cells in all four quadrants of our conceptual figure (Figure S2), but the majority are ‘areas of low concern’ and ‘conservation prioritisation’ (Figure 4; Table S3). This confirms earlier analyses of the lack of importance of high seas fisheries (*20*–*23*), and although the high seas are dominated by areas of low concern, it has vast amounts that fall into ‘conservation prioritisation’ with very few cells in fishery prioritisation or fishery competition. Therefore, a relevant question may be reframed from which parts of the high seas should we protect, to which parts of the high seas should remain as fishing areas (*24*) if any?

**Fig. 4.**
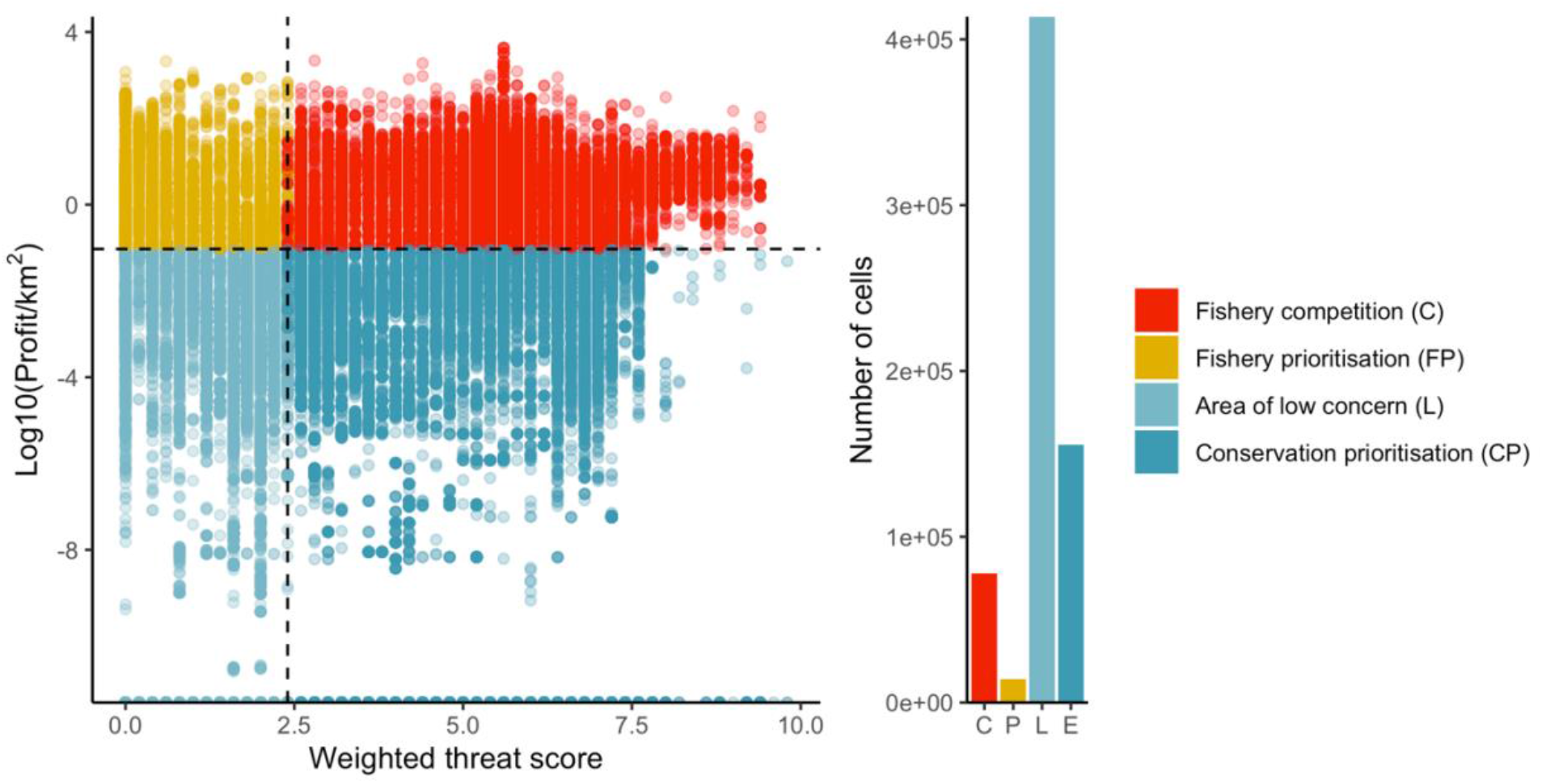
Scatterplot of cells within the High Seas (areas beyond national jurisdiction) according to their categories from our conceptual diagram. Each point is the values for a specific gear type in a 0.5° by 0.5° cell. Dashed lines indicated median values of all cells (High Seas and EEZs).

## Discussion

The United Nations has called for countries to work towards a wide range of Sustainable Development Goals (SDGs). In order to achieve SDG 14 (“Life Below Water”), a greater emphasis needs to placed on fisheries management and protecting marine specific of conservation concern. Previous research has focused on potential trade-offs between fisheries and conservation and where we can search for win-win situations in these two objectives. While it has been shown that often there are benefits to reducing fisheries capacity and fishing to both fisheries (*25*) and conservation (*26*), these benefits are not shared evenly. Reductions in fishing capacity often mean reductions in employment (SDG 8) and seafood supply (SDG 2) (*8*) in the short term (*27*). In addition, closing areas to fisheries can force some fishers out with an often unequal distribution of benefits and costs between different sectors (including eco-tourism) and between different fisheries (SDG 10) (*28*, *29*).

The majority of grid cells within the high seas are rated as being of low conservation concern and low fisheries profitability. While there are likely benefits to coastal fisheries from closing the high seas (*20*, *21*), the conservation benefit of this measure for IUCN Red List species is currently low. Most of the high seas is currently categorized as ‘areas of low concern’ for fisheries and conservation. Only a small area of the high seas (4%) is currently categorized as ‘fishery prioritisation’, and another 16% are areas where there are conservation concerns but also valuable fisheries. The number of cells that are categorized to be ‘conservation prioritisation’ dwarfs the number of cells that are important to fisheries (combined number of cells categorized as ‘fishery prioritisation’ or ‘competition’). This may change in the future as fisheries continue to expand offshore. Therefore, if treated as a whole, the benefits to conservation outweigh the benefits to fisheries in the high seas.

This analysis highlights at a broad scale where fisheries or conservation can be prioritized, and where there are competing aims between these areas. Coastal areas are of large importance to fisheries, especially to small-scale fisheries, but coastal areas are also the most biodiverse regions of the marine realm. These areas are generally categorized as ‘competition’ areas. However, this analysis adds to existing MPA discussions that may lead to less contentious implementation of MPAs where certain fisheries can co-exist depending on the MPA goals (species conservation versus resource conservation) and current fisheries threats.

Our study is static and focuses on data reflective of the present situation. We therefore do not account for the future marine spatial planning challenges associated with changing species ranges (*30*), and the response of fisheries to climate change and changing environmental parameters (*31*, *32*) and their economic and human consequences (*33*). The concepts of this study could be incorporated into models that allow for gear substitution to model fisheries adaptation to climate change along a path that reduces impacts on Threatened species.

Our study may underestimate the impact of some fishing gears based on the descriptions of threats and use for each IUCN species. The study is inherently limited to those species included in the IUCN Red List, as well as to those species that have enough information to be included in the analysis (see supplemental materials). For the species that have gear threat information, it is unlikely these threats are biased towards particular fishing methods as the threats generally highlight all known (fisheries) threats. However, there is known to be a systemic bias taxonomically and geographically for conservation research and species assessed by the Red List (*34*). One area where this is not fully accounted for is the impact of bottom trawls, dredges, and other bottom-impacting gear on seafloor habitat (*35*), which is not fully captured in the IUCN assessments (supplemental materials). In addition, as Data Deficient species are given a risk score of 0, we likely underestimate the risk to these species. For example, a quarter of Chondrichthyes (sharks, rays, and chimaeras) species currently categorized as data deficient were predicted to be Threatened (*36*).

## Conclusion

Our results highlight areas of high conservation concern for particular fishing gears, and areas of high overlap between multiple fishing gear threats and multiple species of conservation concern. We also highlight areas with the potential for low-cost fishing closures leading to maximum protection of species negatively affected by these fishing gears. Interestingly, the study suggests that gillnet fisheries represent a greater overall threat than the often criticised bottom trawl fisheries due to the high fisheries profits derived from many bottom trawl catches. This analysis can help inform future conservation planning with areas of low-cost trade-offs in comparison to areas with much higher costs for equal conservation benefits.

## Acknowledgments

*Acknowledgments: Funding*: TC acknowledges funding support of the Social Sciences and Humanities Research Council of Canada and the Four Year Fellowship from the University of British Columbia. DP acknowledges support from the *Sea Around Us*, which is funded by a number of philanthropic foundations. TCT and URS acknowledge support from the OceanCanada Partnership, funded by the Social Sciences and Humanities Research Council of Canada. URS acknowledges additional funding from the ADM Capital Foundation.

## Author contributions

TC led the design and analysis and wrote the original version of the manuscript. TCT helped in the early analysis and editing the original manuscript. VL contributed data and expertise on the cost of fishing database. TC, TCT, VL, DP, and URS helped analyse results and edited the final manuscript to its improved form.

## Data and materials availability

Data obtained from the *Sea Around Us*, BirdLife International, and the IUCN Red List are publicly available. All code used for processing this data and producing the results of this manuscript are available at www.github.com/timcashion/iucnfishingthreats.

## References

1. J. B. C. Jackson, M. X. Kirby, W. H. Berger, K. a Bjorndal, L. W. Botsford, B. J. Bourque, R. H. Bradbury, R. Cooke, J. Erlandson, J. a Estes, T. P. Hughes, S. Kidwell, C. B. Lange, H. S. Lenihan, J. M. Pandolfi, C. H. Peterson, R. S. Steneck, M. J. Tegner, R. R. Warner, Historical overfishing and the recent collapse of coastal ecosystems. Science (New York, N.Y.). 293, 629–37 (2001).

2. D. A. Kroodsma, J. Mayorga, T. Hochberg, N. A. Miller, K. Boerder, F. Ferretti, A. Wilson, B. Bergman, T. D. White, B. A. Block, P. Woods, B. Sullivan, C. Costello, B. Worm, Tracking the global footprint of fisheries. Science (New York, N.Y.). 359, 904–908 (2018).

3. R. O. Amoroso, A. M. Parma, C. R. Pitcher, R. A. Mcconnaughey, S. Jennings, Comment on “Tracking the global footprint of fisheries”. Science. 361, eaat6713 (2018).

4. N. Queiroz, N. E. Humphries, A. Couto, M. Vedor, I. da Costa, A. M. M. Sequeira, G. Mucientes, A. M. Santos, F. J. Abascal, D. L. Abercrombie, K. Abrantes, D. Acuña-Marrero, A. S. Afonso, P. Afonso, D. Anders, G. Araujo, R. Arauz, P. Bach, A. Barnett, D. Bernal, M. L. Berumen, S. B. Lion, N. P. A. Bezerra, A. V. Blaison, B. A. Block, M. E. Bond, R. W. Bradford, C. D. Braun, E. J. Brooks, A. Brooks, J. Brown, B. D. Bruce, M. E. Byrne, S. E. Campana, A. B. Carlisle, D. D. Chapman, T. K. Chapple, J. Chisholm, C. R. Clarke, E. G. Clua, J. E. M. Cochran, E. C. Crochelet, L. Dagorn, R. Daly, D. D. Cortés, T. K. Doyle, M. Drew, C. A. J. Duffy, T. Erikson, E. Espinoza, L. C. Ferreira, F. Ferretti, J. D. Filmalter, G. C. Fischer, R. Fitzpatrick, J. Fontes, F. Forget, M. Fowler, M. P. Francis, A. J. Gallagher, E. Gennari, S. D. Goldsworthy, M. J. Gollock, J. R. Green, J. A. Gustafson, T. L. Guttridge, H. M. Guzman, N. Hammerschlag, L. Harman, F. H. V. Hazin, M. Heard, A. R. Hearn, J. C. Holdsworth, B. J. Holmes, L. A. Howey, M. Hoyos, R. E. Hueter, N. E. Hussey, C. Huveneers, D. T. Irion, D. M. P. Jacoby, O. J. D. Jewell, R. Johnson, L. K. B. Jordan, S. J. Jorgensen, W. Joyce, C. A. K. Daly, J. T. Ketchum, A. P. Klimley, A. A. Kock, P. Koen, F. Ladino, F. O. Lana, J. S. E. Lea, F. Llewellyn, W. S. Lyon, A. MacDonnell, B. C. L. Macena, H. Marshall, J. D. McAllister, R. McAuley, M. A. Meÿer, J. J. Morris, E. R. Nelson, Y. P. Papastamatiou, T. A. Patterson, C. Peñaherrera-Palma, J. G. Pepperell, S. J. Pierce, F. Poisson, L. M. Quintero, A. J. Richardson, P. J. Rogers, C. A. Rohner, D. R. L. Rowat, M. Samoilys, J. M. Semmens, M. Sheaves, G. Shillinger, M. Shivji, S. Singh, G. B. Skomal, M. J. Smale, L. B. Snyders, G. Soler, M. Soria, K. M. Stehfest, J. D. Stevens, S. R. Thorrold, M. T. Tolotti, A. Towner, P. Travassos, J. P. Tyminski, F. Vandeperre, J. J. Vaudo, Y. Y. Watanabe, S. B. Weber, B. M. Wetherbee, T. D. White, S. Williams, P. M. Zárate, R. Harcourt, G. C. Hays, M. G. Meekan, M. Thums, X. Irigoien, V. M. Eguiluz, C. M. Duarte, L. L. Sousa, S. J. Simpson, E. J. Southall, D. W. Sims, Global spatial risk assessment of sharks under the footprint of fisheries. Nature. 4 (2019), doi:10.1038/s41586-019-1444-4.

5. R. L. Lewison, L. B. Crowder, A. J. Read, S. A. Freeman, Understanding impacts of fisheries bycatch on marine megafauna. Trends in Ecology & Evolution. 19, 598–604 (2004).

6. FAO, “The State of World Fisheries and Aquaculture 2018 - Meeting the sustainable development goals” (Rome, 2018), p. 210.

7. C. White, B. S. Halpern, C. V. Kappel, Ecosystem service tradeoff analysis reveals the value of marine spatial planning for multiple ocean uses. Proceedings of the National Academy of Sciences of the United States of America. 109, 4696–4701 (2012).

8. W. W. Cheung, U. R. Sumaila, Trade-offs between conservation and socio-economic objectives in managing a tropical marine ecosystem. Ecological Economics. 66, 193–210 (2008).

9. T. O. McShane, P. D. Hirsch, T. C. Trung, A. N. Songorwa, A. Kinzig, B. Monteferri, D. Mutekanga, H. V. Thang, J. L. Dammert, M. Pulgar-Vidal, M. Welch-Devine, J. Peter Brosius, P. Coppolillo, S. O’Connor, Hard choices: Making trade-offs between biodiversity conservation and human well-being. Biological Conservation. 144, 966–972 (2011).

10. C. J. Klein, C. Steinback, M. Watts, A. J. Scholz, H. P. Possingham, Spatial marine zoning for fisheries and conservation. Frontiers in Ecology and the Environment. 8, 349–353 (2010).

11. FAO, “The State of World Fisheries and Aquaculture 2016: Contributing to Food Security and Nutrition for All” (Food; Agriculture Organization, Rome, Italy, 2016), p. 200.

12. N. J. Beaumont, M. C. Austen, S. C. Mangi, M. Townsend, Economic valuation for the conservation of marine biodiversity. Marine Pollution Bulletin. 56, 386–396 (2008).

13. Millennium Ecosystem Assessment, Ecosystems and human well-being: Synthesis (Island Press, Washington, DC, 2005).

14. IUCN, IUCN Red List of Threatened Species. Version 2019-2 (2019), (available at www.iucnredlist.org).

15. D. Pauly, D. Zeller, Eds., Sea Around Us: Concepts, Design and Data (2015; http://www.seaaroundus.org).

16. C. C. O’Hara, J. Carlos Villaseñor-Derbez, G. M. Ralph, B. S. Halpern, Mapping status and conservation of global at-risk marine biodiversity. Conservation Letters (2019), doi:10.1111/conl.12651.

17. H. S. Gordon, The Economic Theory of a Common-Property Resource: The Fishery. Journal of Political Economy. 62, 124–142 (1954).

18. T. Cashion, D. Al-abdulrazzak, D. Belhabib, B. Derrick, E. Divovich, D. K. Moutopoulos, S.-l. Noël, M.-L. Palomares, L. C. L. Teh, D. Zeller, D. Pauly, Reconstructing global marine fishing gear use: Catches and landed values by gear type and sector. Fisheries Research. 206, 57–64 (2018).

19. R. Chuenpagdee, L. E. Morgan, S. M. Maxwell, E. A. Norse, D. Pauly, Shifting gears: Assessing collateral impacts of fishing methods in US waters. Frontiers in Ecology and the Environment. 1, 517–524 (2003).

20. U. R. Sumaila, V. W. Lam, D. D. Miller, L. Teh, R. A. Watson, D. Zeller, W. W. Cheung, I. M. Côté, A. D. Rogers, C. Roberts, E. Sala, D. Pauly, Winners and losers in a world where the high seas is closed to fishing. Scientific Reports. 5, 8481 (2015).

21. C. White, C. Costello, Close the High Seas to Fishing? PLoS Biology. 12, 1–5 (2014).

22. L. Schiller, M. Bailey, J. Jacquet, E. Sala, “High seas fisheries play a negligible role in addressing global food security” (2018), pp. 8351–8359.

23. U. R. Sumaila, D. Zeller, R. Watson, J. Alder, D. Pauly, Potential costs and benefits of marine reserves in the high seas. Marine Ecology Progress Series. 345, 305–310 (2007).

24. C. Walters, in Reinventing fisheries management, T. J. Pitcher, D. Pauly, P. Hart, Eds. (Dordrecht, 1998), pp. 279–288.

25. U. R. Sumaila, W. W. Cheung, A. Dyck, K. Gueye, L. Huang, V. W. Lam, D. Pauly, T. Srinivasan, W. Swartz, R. Watson, D. Zeller, Benefits of rebuilding global marine fisheries outweigh costs. PLoS ONE. 7 (2012), doi:10.1371/journal.pone.0040542.

26. M. G. Burgess, G. R. McDermott, B. Owashi, L. E. Peavey Reeves, T. Clavelle, D. Ovando, B. P. Wallace, R. L. Lewison, S. D. Gaines, C. Costello, Protecting marine mammals, turtles, and birds by rebuilding global fisheries. Science (New York, N.Y.). 359, 1255–1258 (2018).

27. U. R. Sumaila, Intergenerational cost – benefit analysis and marine ecosystem restoration. Fish and Fisheries. 5, 329–343 (2004).

28. D. A. Gill, S. H. Cheng, L. Glew, E. Aigner, N. J. Bennett, M. B. Mascia, Social Synergies, Tradeoffs, and Equity in Marine Conservation Impacts. Annual Review of Environment and Resources. 44, annurev–environ–110718–032344 (2019).

29. J. E. Cinner, T. Daw, C. Huchery, P. Thoya, A. Wamukota, M. Cedras, C. Abunge, Winners and Losers in Marine Conservation: Fishers’ Displacement and Livelihood Benefits from Marine Reserves. Society and Natural Resources. 27, 994–1005 (2014).

30. W. W. Cheung, V. W. Lam, K. Kearney, D. Pauly, Projecting global marine biodiversity impacts under climate change scenarios. Fish and Fisheries. 10, 235–251 (2009).

31. G. O. Crespo, D. C. Dunn, G. Reygondeau, K. Boerder, B. Worm, W. W. Cheung, D. P. Tittensor, P. N. Halpin, The environmental niche of the global high seas pelagic longline fleet. Science Advances. 4, 1–14 (2018).

32. T. Young, E. C. Fuller, M. M. Provost, K. E. Coleman, K. S. Martin, B. J. McCay, M. L. Pinsky, Adaptation strategies of coastal fishing communities as species shift poleward. ICES Journal of Marine Science. 76, 93–103 (2019).

33. U. R. Sumaila, T. C. Tai, V. W. Lam, W. W. Cheung, M. Bailey, A. M. Cisneros-Montemayor, O. L. Chen, S. S. Gulati, Benefits of the Paris Agreement to ocean life, economies, and people. Science Advances. 5 (2019) (available at http://advances.sciencemag.org/).

34. M. R. Donaldson, N. J. Burnett, D. C. Braun, C. D. Suski, S. G. Hinch, S. J. Cooke, J. T. Kerr, Taxonomic bias and international biodiversity conservation research. Facets. 1, 105–113 (2016).

35. Committee on Ecosystem Effects of Fishing, Effects of Trawling and Dredging on Seafloor Habitat (National Academy Press, Washington, DC, 2002; http://www.nap.edu/catalog/10323).

36. N. K. Dulvy, S. L. Fowler, J. A. Musick, R. D. Cavanagh, P. M. Kyne, L. R. Harrison, J. K. Carlson, L. N. Davidson, S. V. Fordham, M. P. Francis, C. M. Pollock, C. A. Simpfendorfer, G. H. Burgess, K. E. Carpenter, L. J. Compagno, D. A. Ebert, C. Gibson, M. R. Heupel, S. R. Livingstone, J. C. Sanciangco, J. D. Stevens, S. Valenti, W. T. White, Extinction risk and conservation of the world’s sharks and rays. eLife. 3 (2014), doi:10.7554/eLife.00590.

37. BirdLife International, Handbook of the Birds of the World, No Title (2018), (available at http://datazone.birdlife.org/species/requestdis).

38. M. D. Spalding, H. E. Fox, G. R. Allen, N. Davidson, Z. A. Ferdaña, M. Finlayson, B. S. Halpern, M. A. Jorge, A. Lombana, S. A. Lourie, K. D. Martin, E. McManus, J. Molnar, C. A. Recchia, J. Robertson, Marine Ecoregions of the World: A Bioregionalization of Coastal and Shelf Areas. BioScience (2007), doi:10.1641/b570707.

39. D. Zeller, M.-L. Palomares, A. Tavakolie, M. Ang, D. Belhabib, W. W. Cheung, V. W. Lam, E. Sy, G. Tsui, K. Zylich, D. Pauly, Still catching attention: Sea Around Us reconstructed global catch data, their spatial expression and public accessibility. Marine Policy. 70, 145–152 (2016).

40. D. Grémillet, A. Ponchon, M. Paleczny, M.-L. Palomares, V. Karpouzi, D. Pauly, Persisting Worldwide Seabird-Fishery Competition Despite Seabird Community Decline. Current Biology. 28, 4009–4013.e2 (2018).

41. T. C. Tai, T. Cashion, V. W. Lam, W. Swartz, U. R. Sumaila, Ex-vessel fish price database: disaggregating prices for low-priced species from reduction fisheries. Frontiers in Marine Science. 4, 1–10 (2017).

42. M.-y. Lee, Hedonic Pricing of Atlantic Cod: Effects of Size, Freshness, and Gear. Marine Resource Economics. 29 (2014).

43. F. Asche, J. Guillen, The importance of fishing method, gear and origin: The Spanish hake market. Marine Policy. 36, 365–369 (2012).

44. K. E. Mcconnell, I. E. Strand, Hedonic Prices for Fish: Tuna Prices in Hawaii. American Journal of Agricultural Economics. 82, 133–144 (2000).

45. NMFS, Commercial Fisheries - Annual Landings (2017), (available at http://www.st.nmfs.noaa.gov/commercial-fisheries/commercial-landings/annual-landings/index).

46. V. W. Lam, U. R. Sumaila, A. Dyck, D. Pauly, R. Watson, Construction and first applications of a global cost of fishing database. ICES Journal of Marine Science. 68, 1996–2004 (2011).

47. FAO, “Fishery Statistical Collections: Global capture production. (1950-2015). Accessed through FishStatJ software” (UN FAO Fisheries; Aquaculture Department, Rome, 2017).

48. A. Schuhbauer, R. Chuenpagdee, W. W. Cheung, K. Greer, U. R. Sumaila, How subsidies affect the economic viability of small-scale fisheries. Marine Policy. 82, 114–121 (2017).

49. R Core Team, R Development Core Team. 55 (2017), pp. 275–286.

50. H. Wickham, “tidyverse: Easily Install and Load ‘Tidyverse’ Packages.” (2016), (available at https://cran.r-project.org/package=tidyverse).

51. B. Baumer, D. Udwin, R Markdown (2015),, doi:10.1002/wics.1348.

52. C. Fay, Text Mining with R: A Tidy Approach. Journal of Statistical Software (2018), doi:10.18637/jss.v083.b01.

53. B. Auguie, Package ‘ gridExtra’. R CRAN Project (2017).

54. E. Pebesma, Simple Features for R: Standardized Support for Spatial Vector Data. The R Journal (2019), doi:10.32614/rj-2018-009.

55. E. Pebesma, B. Rowlingson, R. Bivand, Package ‘rgdal’. R-CRAN (2012).

56. Huber, W., Carey, V. J., Gentleman, R., Anders, S., Carlson, M., Carvalho, B. S., Bravo, H. C., Davis, S., Gatto, L., Girke, T., Gottardo, R., Hahne, F., Hansen, K. D., Irizarry, R. A., Lawrence, M., Love, M. I., MacDonald, J., Obenchain, V., Ole’s, A. K., Pag‘es, H., Reyes, A., Shannon, P., Smyth, G. K., Tenenbaum, D., Waldron, L., Morgan, M., C. O. Wilke, H. Wickham, R Core Team, cowplot: Streamlined Plot Theme and Plot Annotations for ‘ggplot2’. Nature Methods (2017).

57. K. Ram, H. Wickham, wesanderson: A Wes Anderson Palette Generator (2018), (available at https://cran.r-project.org/package=wesanderson).

58. B. S. Halpern, C. Longo, D. Hardy, K. L. McLeod, J. F. Samhouri, S. K. Katona, K. Kleisner, S. E. Lester, J. Oleary, M. Ranelletti, A. A. Rosenberg, C. Scarborough, E. R. Selig, B. D. Best, D. R. Brumbaugh, F. S. Chapin, L. B. Crowder, K. L. Daly, S. C. Doney, C. Elfes, M. J. Fogarty, S. D. Gaines, K. I. Jacobsen, L. B. Karrer, H. M. Leslie, E. Neeley, D. Pauly, S. Polasky, B. Ris, K. St Martin, G. S. Stone, U. Rashid Sumaila, D. Zeller, An index to assess the health and benefits of the global ocean. Nature. 488, 615–620 (2012).

