## Supplemental Materials for "Low cost conservation: Fishing gear threats to marine species"

**Affiliations:**

### Supplemental Text

As our study was restricted to Red List marine species with spatial information, we attempted to ensure this was representative of broader patterns in Red List species. Generally, the breakdown between IUCN categories appears to be quite similar between all marine species and those included in this study (Figure S3).

### Habitat modification

Of the 2,603 marine species that have trawling ('trawls unspecified') identified as a threat, only 70 have habitat modification also identified as a threat. When reviewing the threat description for these species, many of them are identified as two separate threats. Only 9 of these have 'destructive fishing practices' identified as a threat.

### Supplemental Discussion

It is important to caveat our study knowing that we cannot fully account for the current extent of humanity's use of the oceans and this study has focused on fisheries. Small-scale fisheries are critical to coastal livelihoods across the world and must not be compromised by overly broad conservation initiatives, but instead included when implementing actions to improve aspects of ocean health (e.g. (*58*)).

In addition, while we use catch and fisheries profits as a proxy for importance, these are not adjusted for their importance to the country (i.e., percentage of GDP from fisheries revenues or percentage of animal protein from locally caught fish), nor do they take into account differences in purchasing power parity where $1000 of fisheries profits is much more valuable in poorer countries than in industrialized countries.

While O’Hara et al. (*16*) highlighted areas of conservation need, we identify the areas that would also meet with more or less fisheries overlap and thus competition. In these areas of competition, we assume the placement of marine protected areas (MPAs) would be more contentious due to the reliance on these natural resources. We also highlight areas with a low trade-off between fisheries profits for the most conservation protection where conservation should be prioritized, and that represent potentially easy wins for conservation.

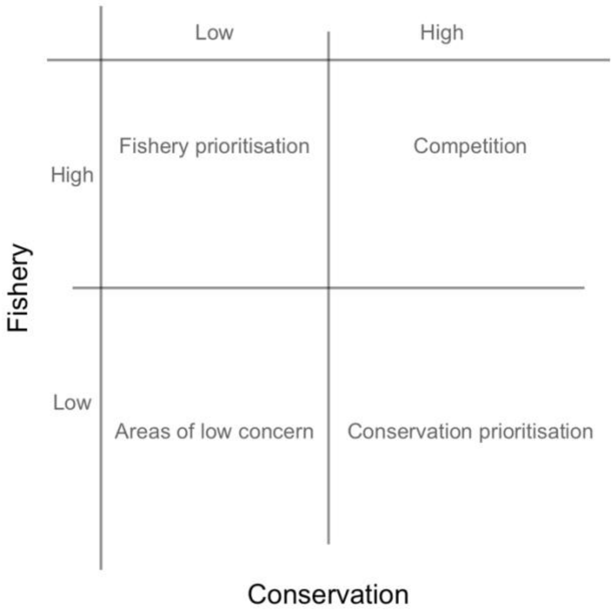


**Fig. S1.** Theoretical basis for competition between fisheries and conservation.

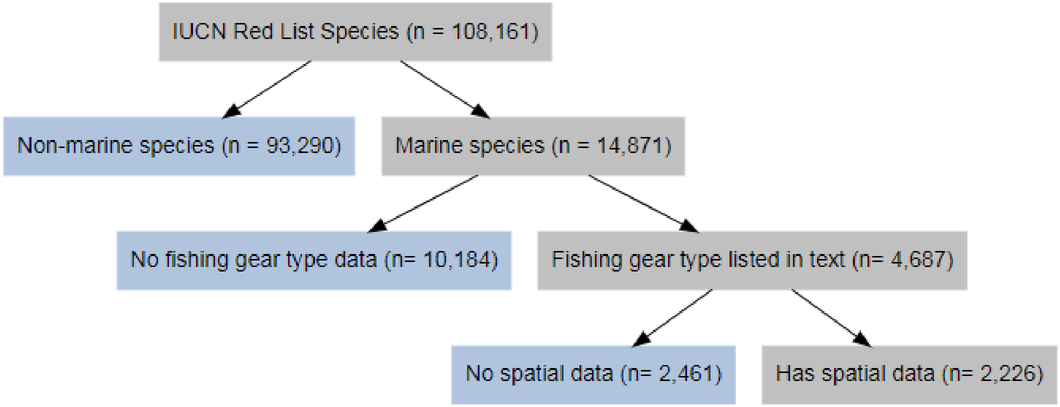


**Fig. S2.** Exclusion reasons for IUCN Red List species for spatial analysis.

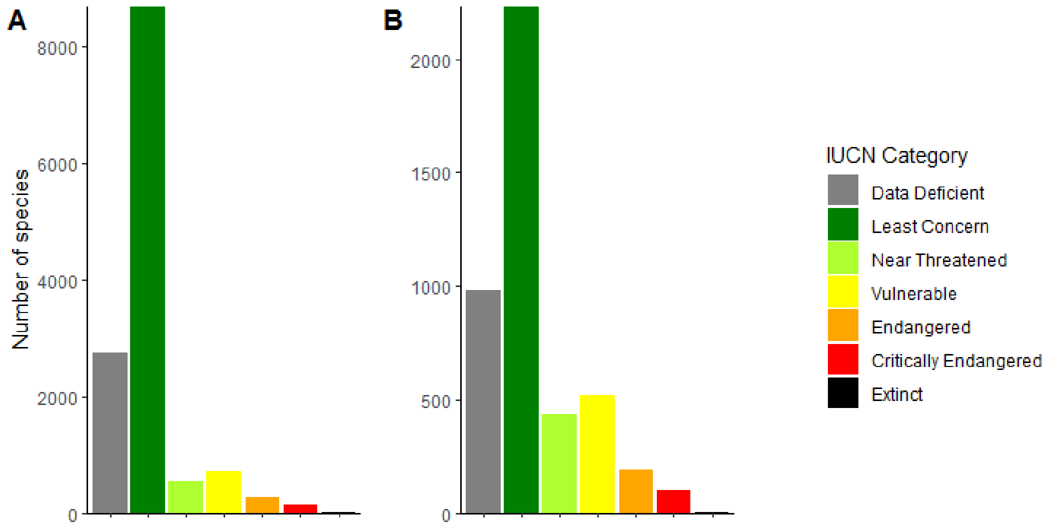


**Fig. S3.** IUCN category for all marine species (A) and marine species included in this study (B).

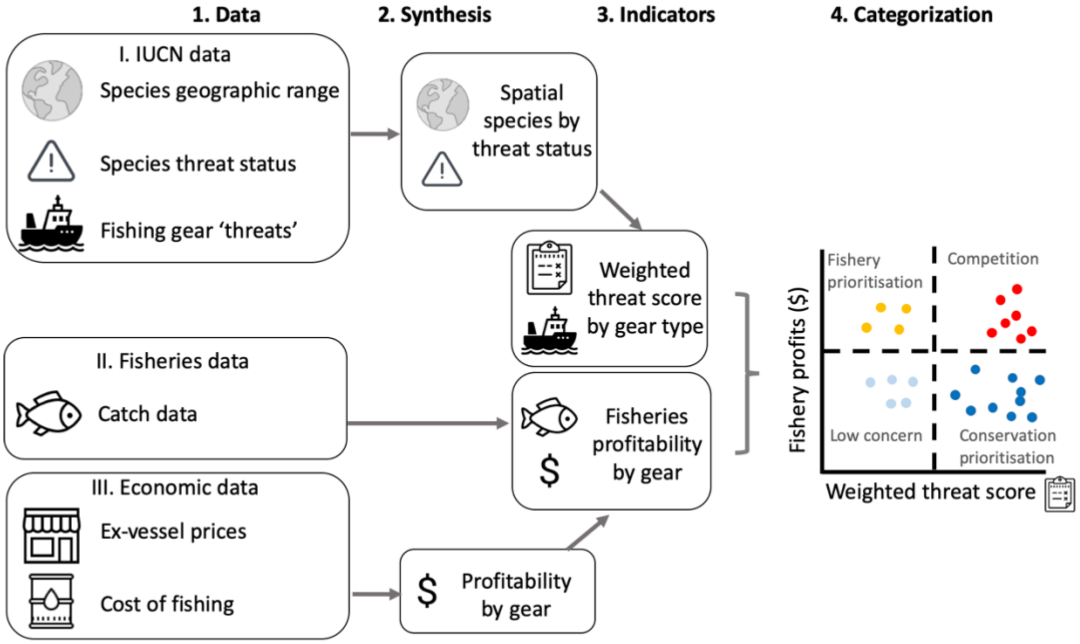


**Fig. S4.** Conceptual methods figure combing I) IUCN data on species geographic range, threat status and fishing gear threats (14), II) Sea Around Us fisheries catch data (15), and III) fisheries ex-vessel prices (41) and cost of fishing data (46).

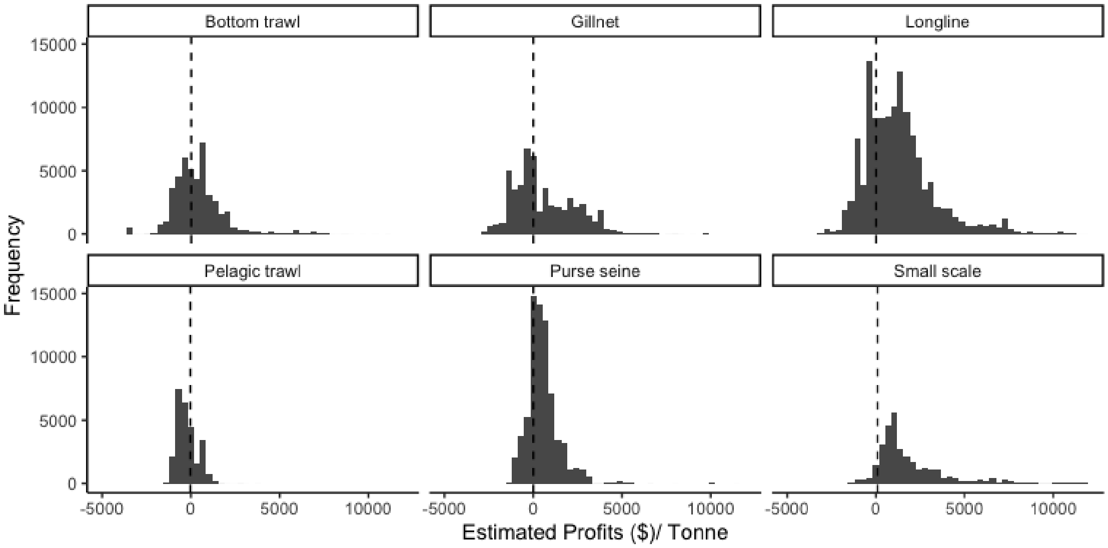


**Fig. S5.** Profit estimated per tonne by gear type where every observation is a fishing entity fishing with a gear in a particular area (0.5° by 0.5° cell). Dashed line indicates mean value for gear type.

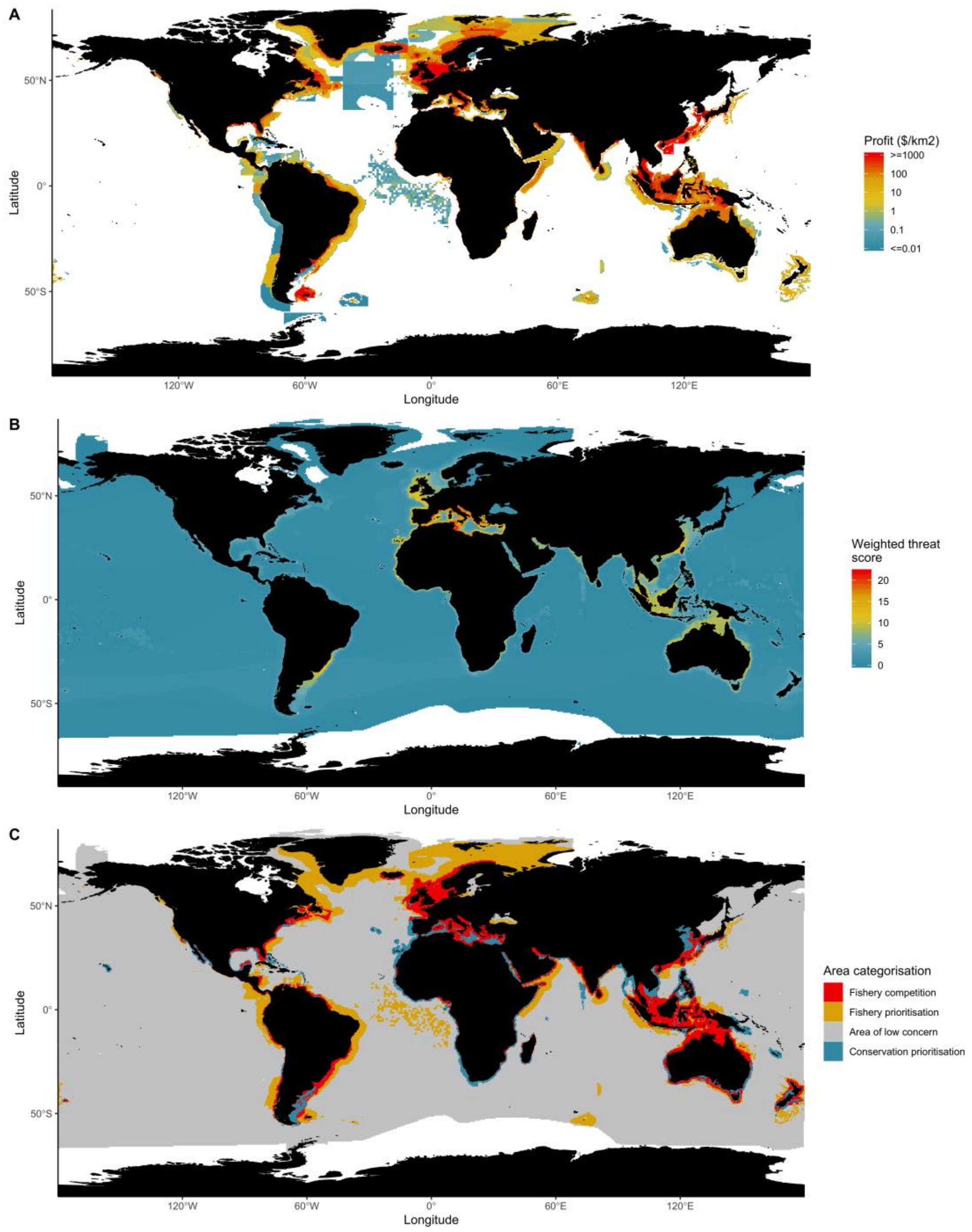


**Fig. S6.** Bottom trawl: Distribution of profits (A), weighted threat score (B), and categorisation (C).

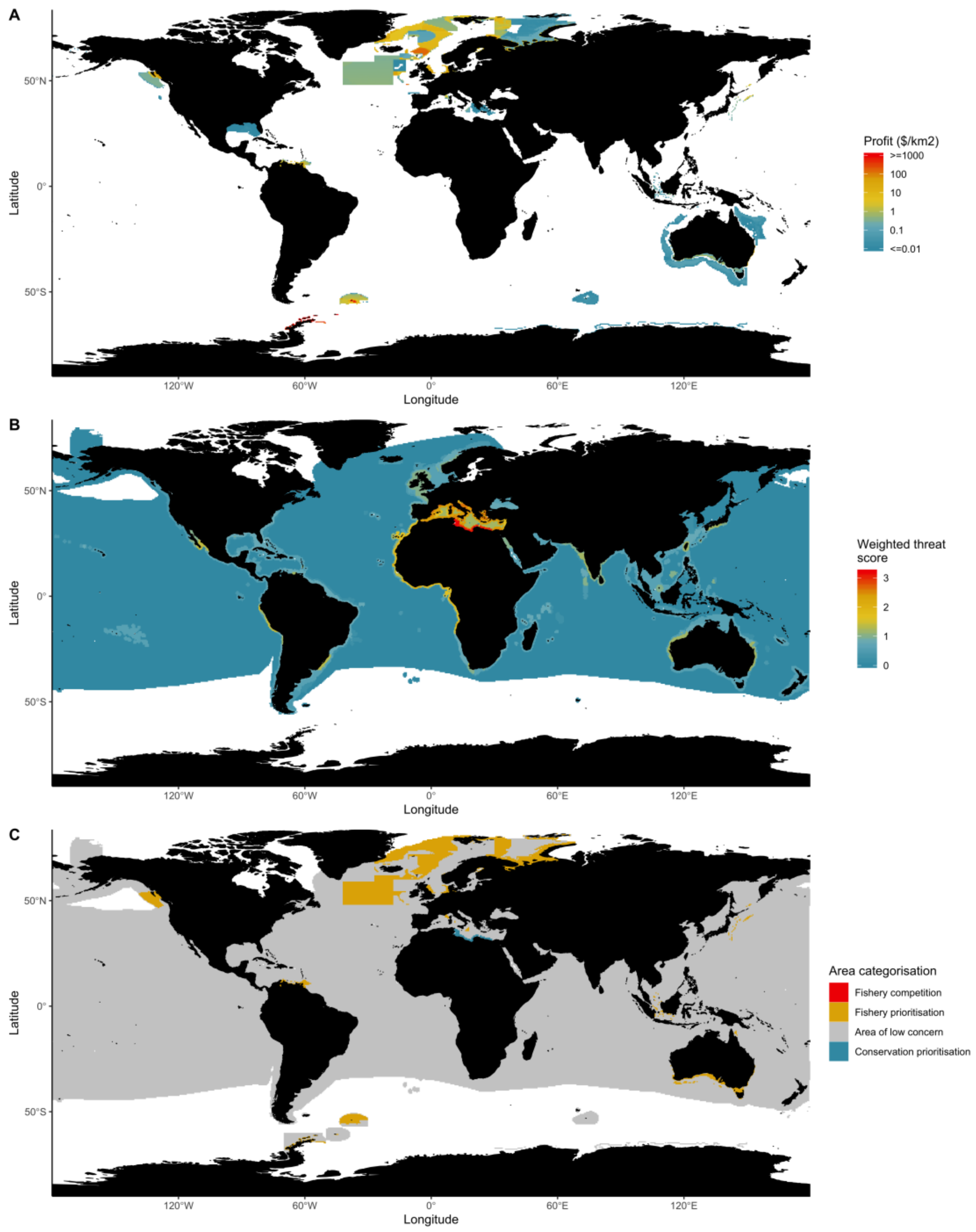


**Fig. S7.** Pelagic trawl: Distribution of profits (A), weighted threat score (B), and categorisation (C).

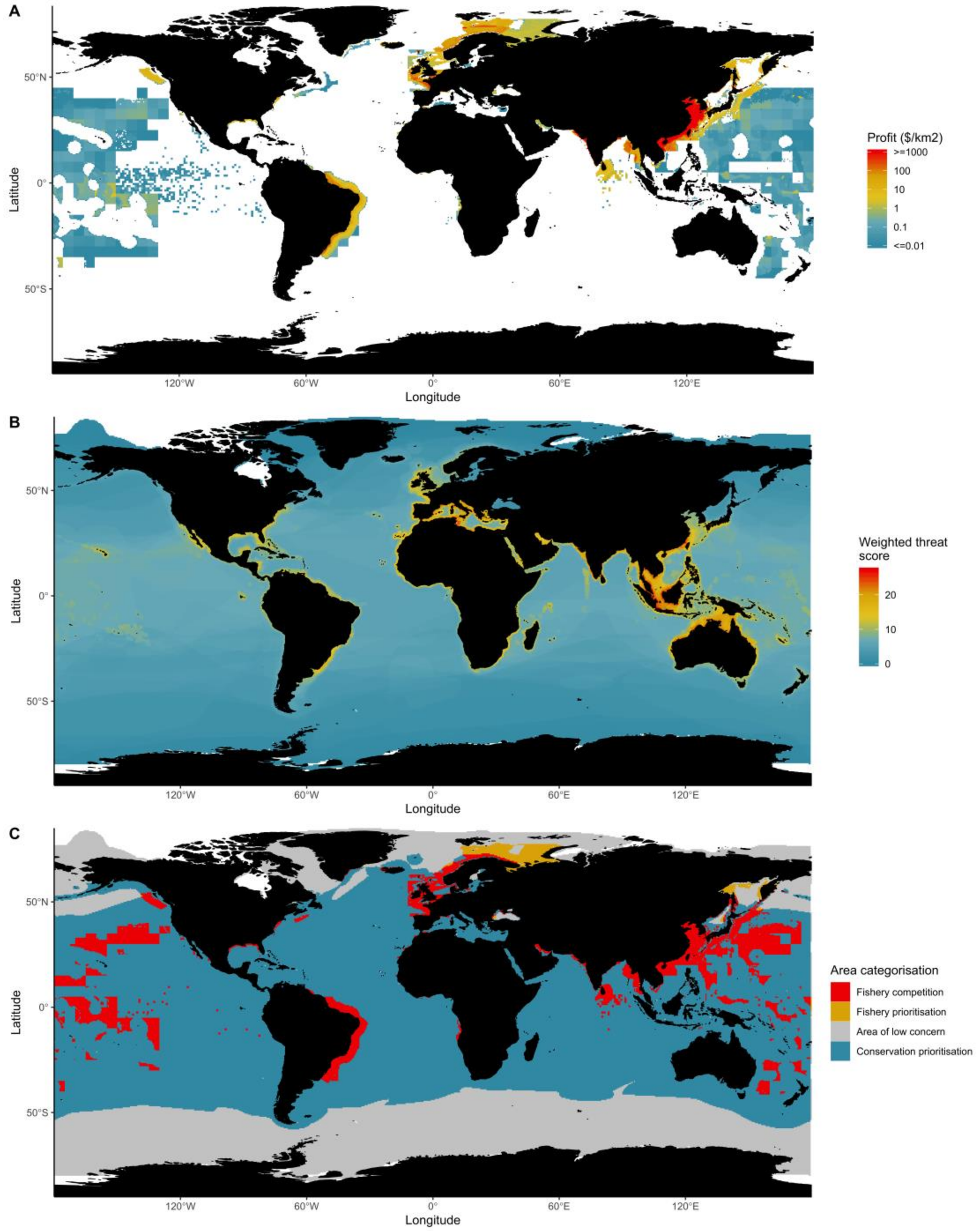


**Fig. S8.** Gillnet: Distribution of profits (A), weighted threat score (B), and categorisation (C).

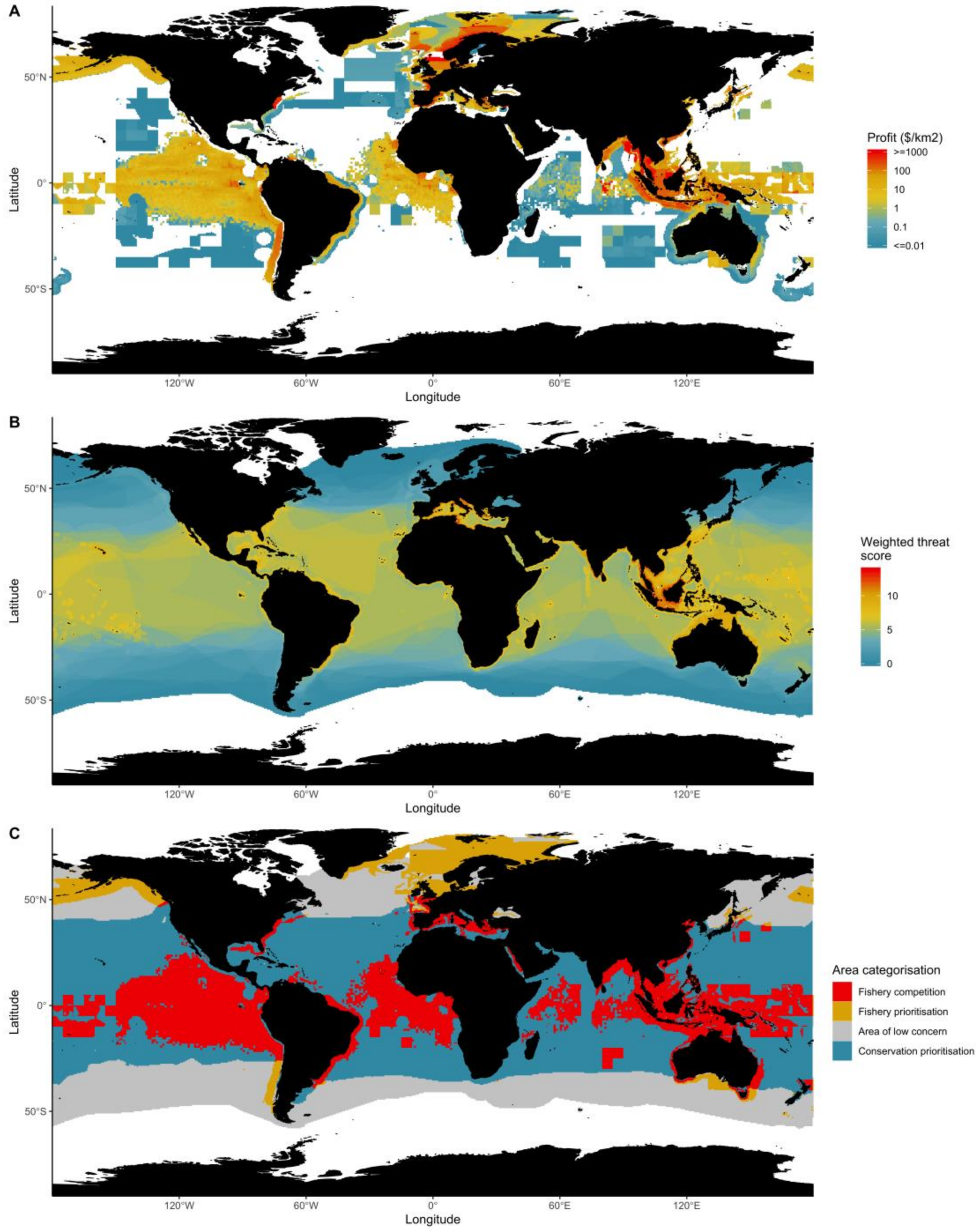


**Fig. S9.** Purse seine: Distribution of profits (A), weighted threat score (B), and categorisation (C).

**Table S1.**

Weighting of IUCN Categories based off (12).

| Former code | Category | Current Code | Category Score |
|---|---|---|---|
| VU | Vulnerable | VU | 0.4 |
| V | Vulnerable | VU | 0.4 |
| CR | Critically Endangered | CR | 0.8 |
| EN | Endangered | EN | 0.6 |
| E | Endangered | EN | 0.6 |
| NT | Near Threatened | NT | 0.2 |
| LC | Least Concern | LC | 0.0 |
| LR/nt | Lower Risk/near threatened | NT | 0.2 |
| LR/lc | Lower Risk/least concern | LC | 0.0 |
| DD | Data Deficient | DD | NA |
| R | Rare | EN | 0.6 |
| O | Out of Danger | LC | 0.0 |
| NA | “Very rare but believed to be stable or increasing” | CR | 0.8 |
| NA | “Status inadequately known-survey required or data sought” | DD | NA |
| LR/cd | Lower Risk/conservation dependent | VU | 0.4 |
| K | Insufficiently Known | DD | NA |
| NA | “Very rare and believed to be decreasing in numbers” | CR | 0.8 |
| NA | “Less rare but believed to be threatened-requires watching” | EN | 0.6 |
| I | Indeterminate | DD | NA |
| EW | Extinct in the Wild | EX | 1.0 |
| EX | Extinct | EX | 1.0 |
| Ex | Extinct | EX | 1.0 |
| NE | Not Evaluated | NE | NA |
| N.E. | Not Evaluated | NE | NA |
| T | Threatened | EN | 0.6 |
| NR | Not Recognized | DD | NA |

**Table S2.**

Search terms used for identifying gear types in IUCN narrative text.

| Gear type | Search terms |
|---|---|
| Bottom trawl | bottom trawl; deepwater trawl; demersal trawl; prawn trawl; benthic trawl; otter trawl; beam trawl; shrimp trawl |
| Pelagic trawl | pelagic trawl; shallow trawl; midwater trawl; surface trawl |
| Trawl | trawl; trawling |
| Longlines | longline; hooks |
| Lines | hook and line; hook and; and line; hook; line; pole and; jig; hand line |
| Nets | net |
| Seines | seine; purse; encircling |
| Gillnets | gill net; gillnet; entangl; entangle |
| Dredges | dredge; drag |
| Traps | pot; trap; fyke |
| Drifting longlines | drifting longlines; drifting lines |
| Squid jiggers | jig; squid jiggers; jiggers; squid jigging; jigging |
| Small scale | small scale; artisanal; subsistence; small-scale |
| Others | harpoon; bagnets; bag; spear |

**Table S3.**

Area of high seas by categorization.

| Category | Water Area (million $km^2$) | Proportion of High Seas (%) |
|---|---|---|
| Fishery competition | 453 | 16.0 |
| Fishery prioritization | 112 | 3.9 |
| Area of low concern | 1467 | 51.8 |
| Conservation prioritization | 802 | 28.3 |

**Table S4.**

Gear types identified in the *Sea Around Us* database.

| Gear type | Super code |
|---|---|
| Bottom trawl | Bottom trawl |
| Shrimp trawl | Bottom trawl |
| Beam trawl | Bottom trawl |
| Otter trawl | Bottom trawl |
| Pelagic trawl | Pelagic trawl |
| Lines | Longline |
| Pole and line | Longline |
| Longline | Longline |
| Hand lines | Longline |
| Encircling nets | Purse seine |
| Purse seine | Purse seine |
| Small encircling nets | Purse seine |
| Gillnet | Gillnet |
| Trammel nets | Gillnet |
| Other | Other |
| Pots or traps | Other |
| Other nets | Other |
| Dredge | Other |
| Hand or tools | Other |
| Dragged gear | Other |
| Mixed gear | Other |
| Artisanal fishing gear | Small scale |
| Long distance small scale | Small scale |
| Recreational fishing gear | Small scale |
| Subsistence fishing gear | Small scale |
| Cast nets | Small scale |
| Bagnets | Small scale |
| Harpoon | Small scale |
| Hand or tools | Small scale |
| Small scale seine nets | Small scale |
| Small scale encircling nets | Small scale |
| Small scale trammel net | Small scale |
| Small scale gillnets | Small scale |
| Small scale other nets | Small scale |

| | |
|---|---|
| Small scale pots or traps | Small scale |
| Small scale lines | Small scale |
| Small scale hand lines | Small scale |
| Small scale pole lines | Small scale |
| Small scale longline | Small scale |
| Small scale purse seine | Small scale |
| Unknown class | Unknown |
| Unknown by source | Unknown |
| Unknown by author | Unknown |

**Table S5.**

Regression results for gear types used and their gear multipliers used.

| Gear type | Estimate | Std.error | T-Statistic | P.value | Multiplier_mean | Multiplier_ll | Multiplier_ul |
|---|---|---|---|---|---|---|---|
| Gillnet | -0.046 | 0.017 | -2.616 | *** | 0.955 | 0.921 | 0.990 |
| Longline | 0.267 | 0.016 | 16.785 | *** | 1.306 | 1.275 | 1.338 |
| Other | 0.055 | 0.015 | 3.735 | *** | 1.057 | 1.028 | 1.086 |
| Pelagic trawl | -0.620 | 0.067 | -9.304 | *** | 0.538 | 0.407 | 0.669 |
| Pots and traps | 0.116 | 0.017 | 7.016 | *** | 1.123 | 1.091 | 1.156 |
| Purse seine | -0.196 | 0.022 | -8.774 | *** | 0.822 | 0.778 | 0.866 |
| Small scale | 0.295 | 0.020 | 14.777 | *** | 1.343 | 1.304 | 1.382 |
| Bottom trawl | 0.000 | 0.000 | 0.000 | Not Estimated | 1.000 | 1.000 | 1.000 |
| Unknown | 0.000 | 0.000 | 0.000 | Not Estimated | 1.000 | 1.000 | 1.000 |
